# Primary sex determination in chickens depends on DMRT1 dosage, but gonadal sex does not determine secondary sexual characteristics in adult birds

**DOI:** 10.1101/2020.09.18.303040

**Authors:** Jason Ioannidis, Gunes Taylor, Debiao Zhao, Long Liu, Alewo Idoko-Akoh, Daoqing Gong, Robin Lovell-Badge, Silvana Guioli, Mike McGrew, Michael Clinton

## Abstract

In birds, males are the homogametic sex (ZZ) and females the heterogametic sex (ZW), and primary sex determination is thought to depend on a sex chromosome gene dosage mechanism. Previous studies have suggested that the most likely sex-determinant is the Z chromosome gene *DMRT1* (Doublesex and Mab-3 Related Transcription factor 1). To clarify this issue, we used a CRISPR-Cas9 based mono-allelic targeting approach and sterile surrogate hosts to generate birds with targeted mutations in the *DMRT1* gene. The resulting chromosomally male (ZZ) chicken with a single functional copy of *DMRT1* developed ovaries in place of testes, demonstrating the avian sex determining mechanism is based on DMRT1 dosage. These ZZ ovaries expressed typical female markers and showed clear evidence of follicular development. However, these ZZ adult birds with an ovary in place of testes were indistinguishable in appearance to wild type adult males, supporting the concept of cell-autonomous sex identity (CASI) in birds. In experiments where oestrogen synthesis was blocked in control ZW embryos, the resulting gonads developed as testes. In contrast, if oestrogen synthesis was blocked in ZW embryos that lacked *DMRT1*, the gonads invariably adopted an ovarian fate. Our analysis shows that DMRT1 is the key sex determination switch in birds and that it is essential for testis development, but that production of oestrogen is also a key factor in primary sex determination in chickens, and that this production is linked to DMRT1 expression.

## Introduction

Primary sex determination is the process whereby the developing gonad differentiates into either a testis or an ovary. In general, the genetic factors that regulate gonadal sex differentiation in vertebrates are well conserved, although the mechanisms that initiate the process, and the hierarchical interactions of the factors involved, can vary considerably between species. Key conserved male differentiation factors include DMRT1 (Doublesex and Mab-3 Related Transcription factor 1) and AMH (anti-Mullerian hormone), although these are utilised in different ways in different species^1^. For example, fishes employ a variety of sex-determining genes, including *dmrt1*, dmrt1y (Y-linked *DMRT1*), *sdy* (sexually dimorphic on Y-chromosome), *amhy* (Y-linked AMH) and *amhr2* (AMH receptor type-2). *Dmrt1* homologs and paralogs, such as *dmw* (W-linked *DMRT1*), are also utilised by some amphibians and reptiles, and sometimes under the control of external stimuli^2-5^. Although DMRT1 does not drive primary sex determination in mice and humans, it does play a role of maintaining male somatic cell sex identity in adult testes^1^. Factors that play key roles in gonadal female sex determination in many vertebrates are FOXL2 (Forkhead box L2) and oestrogen signalling (E_2_). For example, in Tilapia, a Foxl2/Dmrt1 balance appears to control sexual differentiation by regulating E_2_ production through aromatase expression^6^. While E_2_ is not a primary sex-determining factor in most mammals, it is able to override genetic sex determination (GSD) in marsupial neonates^7^. In chickens, blocking E_2_ synthesis in female embryos leads to masculinisation of the gonads, while the addition of E_2_ to male embryos leads to feminisation of the gonads^8-10^.

In birds, the male is the homogametic sex (ZZ) and the female is the heterogametic sex (ZW), but, as yet, there is no evidence for an ovary-determining gene located on the female-specific W-chromosome^11^. It is widely accepted that primary sex-determination in birds is likely to depend on a gene dosage mechanism based on a Z chromosome gene(s)^11^. The most likely candidate gene is the Z chromosome gene *DMRT1*^12^; *DMRT1* expression is restricted to cells of the gonads and the Mullerian ducts and it is expressed at higher levels in the male than in the female at the time of sex determination^13,14^. *In ovo* manipulation studies show that a reduction in DMRT1 levels leads to feminisation of the genetically male (ZZ) gonad^15^ and that overexpression of *DMRT1* leads to masculinisation of the genetically female (ZW) gonad^16^.

To elucidate the role of DMRT1 dosage in chicken sex determination, we used a novel, efficient CRISPR-Cas9 targeting approach and surrogate germ cell hosts to generate chickens with targeted mutations in *DMRT1* and analysed the effects on gonadal development. Here, we clearly demonstrate that avian gonadal sex fate is dependent on *DMRT1* dosage, and that the mechanism involves moderation of E_2_ production. Presence of DMRT1 is essential for testicular differentiation, but not for the early stages of ovarian differentiation. Our analysis further supports the concept of cell-autonomous sex identity (CASI)^17^, as our results show the development of secondary sexual characteristics of non-reproductive tissues in birds is independent of gonadal sex.

## Results

### Generation of *DMRT1*-mutant birds using surrogate hosts

To generate *DMRT1* knockout chickens we used CRISPR-Cas9 to target the *DMRT1* gene in cultured chicken primordial germ cells (PGCs). As DMRT1 is essential for meiosis and gametogenesis in mammals^18,19^, we targeted a loss of function mutation into a single *DMRT1* allele in ZZ PGCs^20^. ZZ germ cells heterozygous for loss-of-function mutations in essential meiotic genes will successfully navigate meiosis and produce functional gametes^21^. We simultaneously delivered a high fidelity CRISPR/Cas9 vector and two ssDNA oligonucleotides into *in vitro* propagated male tdtomato^+^ heterozygote PGCs: one oligonucleotide to create a premature stop codon and a PAM mutation, and a second oligonucleotide, which contained a PAM mutation encoding a synonymous amino acid change in *DMRT1* (Supplementary Table 1). We isolated clonal male PGC populations and identified clones containing the correct (ZZ *DMRT1*^+/-^; formatted as Z^D+^Z^D-^ for simplicity hereafter) mutations in the *DMRT1* locus (n = 10 of 25 clones) (Figure 1a, Supplementary Figure 1 and Methods).

**Figure 1.**
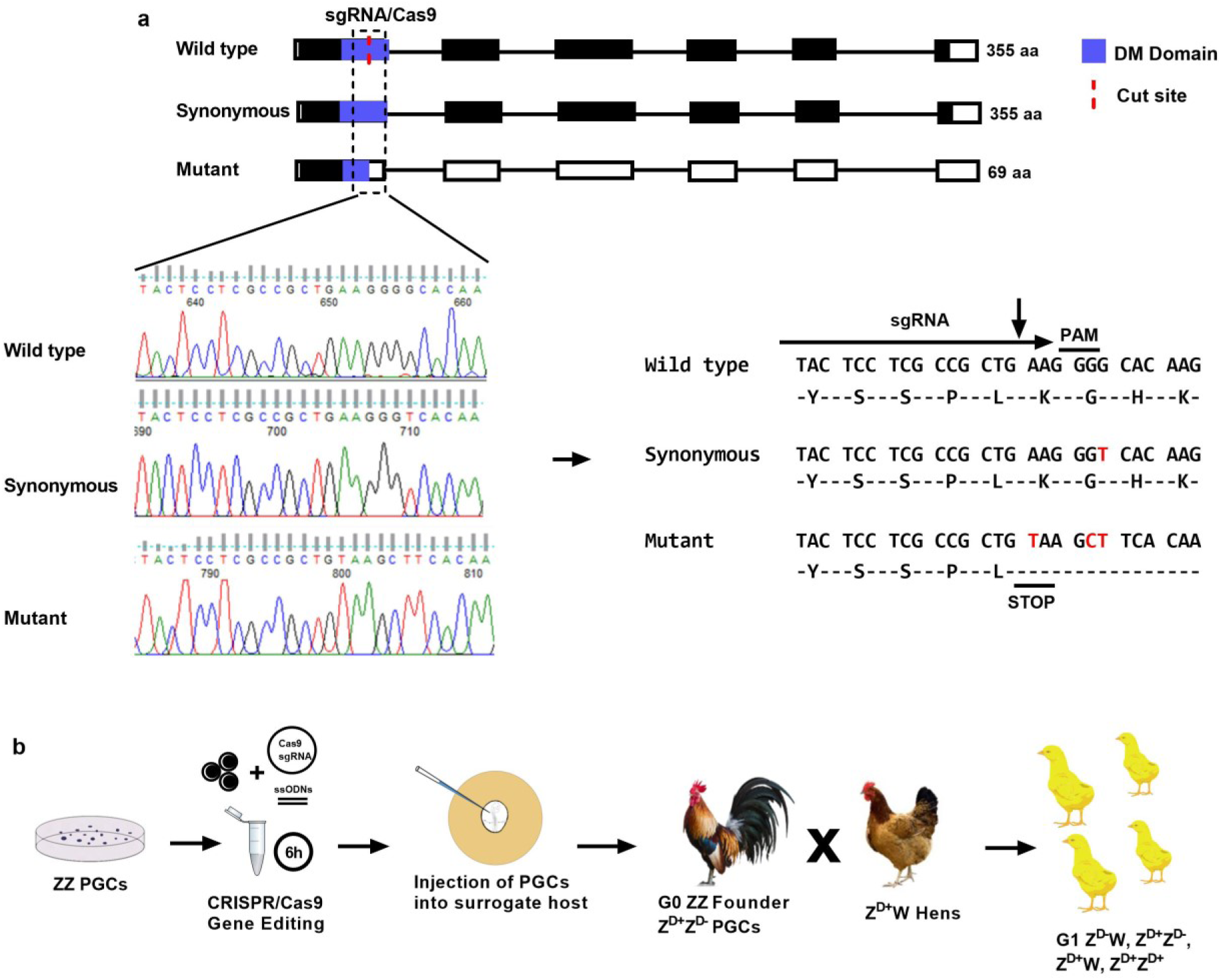
Genome editing of *DMRT1* mutations and genetic crosses. a) Diagram of the *DMRT1* locus in ZZ wild type and edited ZZ PGC clones carrying a synonymous mutation and a loss of function mutation. Details of the Sanger sequencing traces and resulting nucleotide sequences are shown. The non-synonymous change introduced in one allele generates a stop-codon and a frame-shift in the sequence, resulting in a predicted 69 aa truncated protein, which lacks part of the DNA binding domain b) Diagram illustrating the overall technical approach and the mating used to produce *DMRT1-*mutant offspring.

Targeted (Z^D+^Z^D-^) PGCs were injected into transgenic surrogate host chicken embryos containing an inducible Caspase9 targeted to the germ cell-specific *DAZL* locus (Ballantyne et al, under review). Treatment of iCaspase9 host embryos with the dimerization drug, AP20187 (B/B) ablates the endogenous germ cells, such that the only gametes that develop are derived from donor PGCs. The surrogate host (G_0_ founder) chicks were hatched, raised to sexual maturity and then surrogate (G_0_) males (Z^D+^Z^D-^) were naturally mated to Z^D+^W wild type hens (Figure 1b). This mating produced chromosomally male and female G_1_ offspring that were wild type for *DMRT1* (Z^D+^Z^D+^ and Z^D+^W), chromosomally male birds that were heterozygous for functional *DMRT1* (Z^D+^Z^D-^) and chromosomally female birds that lacked functional *DMRT1* (Z^D-^W). PCR and RFP fluorescence expression indicated that 51.6 % of DMRT1 embryos were RFP-positive, suggesting that all offspring derived from exogenous PGCs (see Methods and Supplementary Table 3 for *DMRT1*-allele transmission data).

### ZZ *DMRT1* heterozygote embryos show gonadal sex reversal

Fertile G_1_ eggs from G_0_ founder males mated to wild type females were incubated and examined for gonadal development. Our initial characterisations were performed on embryos at day 13.5 of development (E13.5), as clear morphological differences between male and female gonads are apparent by this stage. As expected, in E13.5 ZZ chick embryos, the testes appeared as two similar sized, cylindrical structures lying on either side of the midline, while ZW embryos contained a left ovary, which acquired an elongated flattened appearance and a small right ovary, which subsequently regressed. The E13.5 testis comprised a core medulla containing germ cell-filled sex cords, while the left ovary contained a relatively unstructured medulla surrounded by a thickened cortex containing germ cells (Figure 2a).

**Figure 2.**
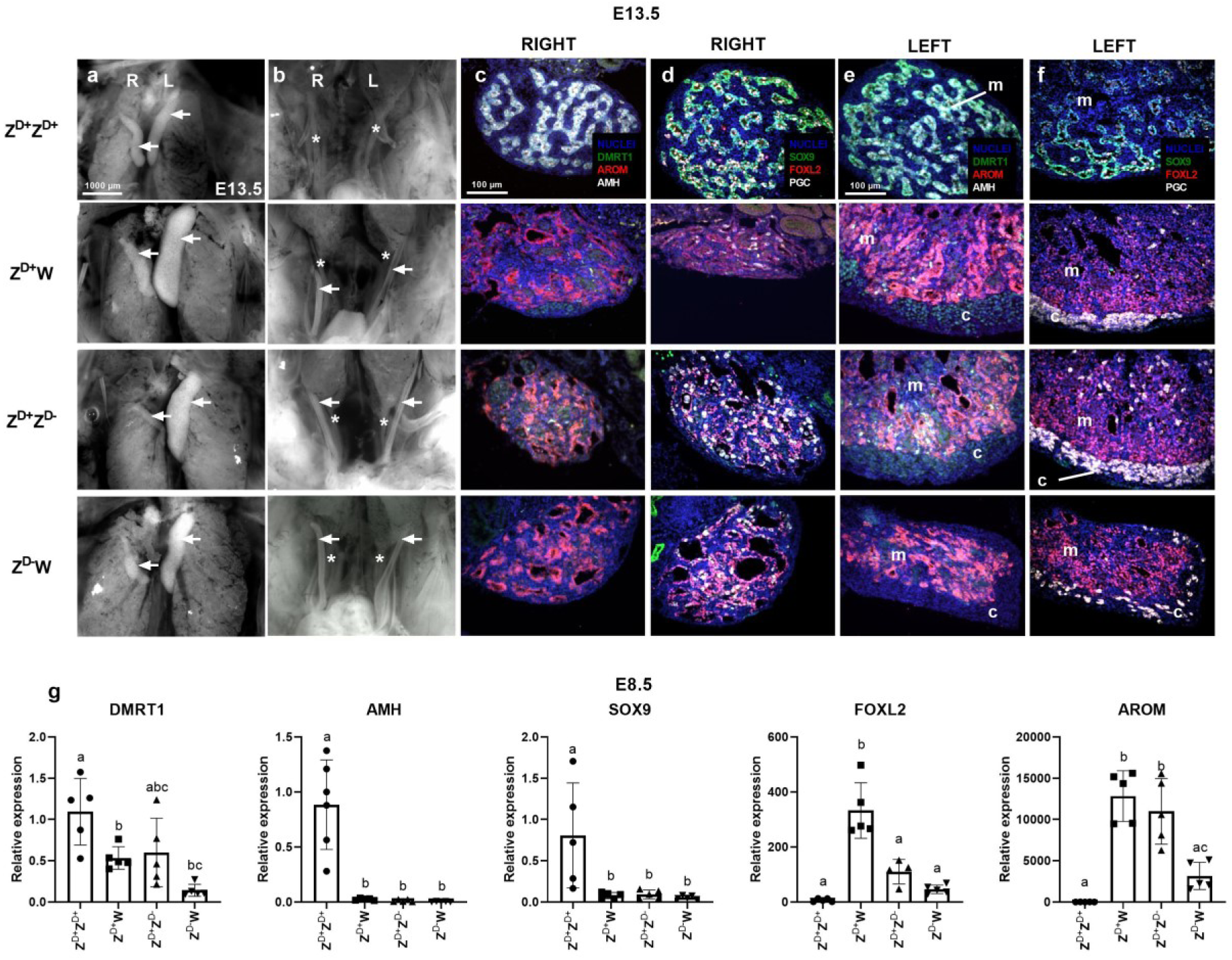
Gonadal development in *DMRT1*-mutant embryos. Gross morphology of gonads (a) and Mullerian ducts (b) in Z^D+^Z^D+^ and Z^D+^W embryos and Z^D+^Z^D-^and Z^D-^W *DMRT1*-mutant embryos (n = 3-7 embryos per genotype). Immuno-sections from right and left gonads from E13.5 wild type and *DMRT1-*mutant embryos (c-f). Expression of DMRT1, aromatase (AROM) and AMH (c, e) and SOX9, FOXL2 and of PGC-specific marker (VASA) (d,f). A minimum of three embryos of each genotype were examined. Arrows indicate gonads in (a) and Mullerian ducts in (b). Asterisks indicate Wolffian ducts in (b). c=cortex; m=medulla. (g) Relative gene expression of *DMRT1* and of testis and ovary markers in gonads of E8.5 wild type and *DMRT1-*mutant embryos. Individual expression levels were calculated relative to levels in Z^D+^Z^D+^. Five replicates on pools of two gonads per genotype. Bars represent mean ± standard deviation. Different letters specify statistically significant groups, P < 0.05.

Examination of the gross morphology of the gonads in Z^D+^Z^D-^ embryos, however, showed that the targeted mutation of *DMRT1* had a significant effect on gonadal development with clear morphological signs of sex reversal (Figure 2a). Unlike the typical paired structures seen in the wild type ZZ embryo, the Z^D+^Z^D-^ clearly contained an ovary-sized structure on the left side and a much smaller structure on the right side, like the Z^D+^W control (n = 5 of 5). In Z^D-^W embryos, the left gonad also appeared to be an ovary, although smaller in size than the wild type counterpart (n = 3 of 3; Figure 2a).

It is interesting to note that, by E13.5, both Mullerian ducts had regressed in the Z^D+^Z^D+^ male, while both Mullerian ducts were retained in Z^D-^W embryos, similar to Z^D+^W embryos (Figure 2b). This result is unexpected, as it was previously published that downregulation of DMRT1 blocks Mullerian duct formation^22^. We also observed that the right Mullerian ducts of both Z^D-^W and to Z^D+^W embryos showed early signs of regression, while, in contrast, the right Mullerian duct of Z^D+^Z^D-^ embryos showed no sign of regression (Figure 2b).

Sections of E13.5 gonads were examined by immunohistochemistry (IHC) to reveal spatial expression patterns of DMRT1 and of established testis (AMH, SOX9 [SRY-box 9]) and ovary (FOXL2, aromatase [CYP19A1-Cytochrome P450 Family 19 Subfamily A member 1]) marker proteins, and PGC-specific markers (Figure 2 c-f). Sections from both right and left Z^D+^Z^D+^ gonads showed a typical male medulla with obvious sex cords comprised of PGCs and somatic cells that expressed DMRT1, SOX9 and AMH, overlaid by a thin epithelial layer. In contrast, the right and left Z^D+^W gonads were structurally distinct. As expected, the medulla of both right and left gonads expressed FOXL2 and aromatase; however, the right gonad was markedly smaller in size. In addition, the left gonad was enclosed within an obvious thickened cortex on the ventral surface, which contained the PGCs. Analyses of sections of gonads from Z^D+^Z^D-^ embryos revealed that they were indistinguishable from Z^D+^W ovaries in terms of structure and molecular profiles. The medullary regions expressed FOXL2 and aromatase and did not contain sex cords or express SOX9 or AMH. DMRT1 was expressed at low levels and the left medulla was surrounded by a PGC-containing cortex typical of a Z^D+^W ovary. In Z^D-^W embryos, both gonads were reduced in size compared to Z^D+^W gonads, but otherwise appeared to be typical ovaries; left and right medullas were FOXL2- and aromatase-positive, and SOX9- and AMH-negative, and the left gonad included a PGC-containing cortex. It is clear from this analysis that the loss of a single functional copy of *DMRT1* leads to ZZ gonadal sex-reversal in chickens.

Similar analyses were performed on embryos collected at E5.5, E6.5 and E8.5 (Supplementary Figure 2a-h). At all stages the gonads of the Z^D+^Z^D-^ embryos, resembled those of wild type ZW embryos rather than wild type ZZ embryos and exhibited testis to ovary sex-reversal. The gonads of Z^D-^W embryos were reduced in size compared to those of wild type Z^D+^W embryos at these stages, but otherwise exhibited structural and functional development typical of ovaries. However, we did observe a slight delay in the upregulation of aromatase in Z^D-^W gonads compared to both Z^D+^Z^D-^ and Z^D+^W embryos (Supplementary Figure 2b).

To confirm that the introduction of a stop codon into the *DMRT1* locus reduces DMRT1 protein levels in heterozygote and homozygote animals, protein extracts from embryonic stage E8.5 gonads were subjected to a Western blot analysis. We observed a reduction in DMRT1 protein levels in Z^D+^Z^D-^ sex-reversed gonads compared to Z^D+^Z^D+^ testes, to levels similar to that in Z^D+^W ovaries. A complete loss of DMRT1 protein was observed in Z^D-^W gonads (Supplementary Figure 3a).

To quantitate the expression of individual gonadal genes, qPCR was performed on RNA extracted from E6.5 and E8.5 gonads. We compared relative expression of *DMRT1* and of testis (*SOX9, AMH*) and ovary (*FOXL2*, aromatase) specific markers in all four genotypes studied. Expression levels at E8.5 relative to expression in Z^D+^Z^D+^ gonads are shown in figure 2g (E6.5 profiles are shown in Supplementary Figure 3b). As expected, the expression levels of *DMRT1* in Z^D+^Z^D+^ gonads were approximately twice that seen in Z^D+^W gonads, while the levels in the latter and in Z^D+^Z^D-^ gonads were similar. Low levels of mutated *DMRT1* transcripts were detected in gonads of Z^D-^W embryos that purportedly lack full-length DMRT1 protein. Relative to Z^D+^Z^D+^ gonads, expression of the ‘male’ markers *SOX9* and *AMH* was essentially absent in Z^D+^Z^D-^ sex reversed gonads and equivalent to levels in control Z^D+^W ovaries. In contrast, there was significant expression of the ‘female’ marker *FOXL2* in Z^D+^Z^D-^ gonads. Although *FOXL2* transcript levels in the latter were lower than those in wild type ovaries, IHC analyses suggested that FOXL2 protein levels were similar (Figure 2c).

Expression levels of aromatase in Z^D+^Z^D-^ gonads were similar to those found in control Z^D+^W ovaries. Expression patterns typical of ovaries were also evident in gonads from Z^D-^W embryos completely lacking DMRT1, although the levels of ovary-specific markers were reduced compared to both Z^D+^W and Z^D+^Z^D-^ gonads.

It is clear from these analyses that gonadal development in Z^D+^Z^D-^ embryos is similar to that seen in control ovaries of ZW female embryos.

### Meiosis in *DMRT1*-mutant embryos

*DMRT1* is also highly expressed in germ cells and has been implicated in the control of meiotic entry and progression in different vertebrate species^18,23^. To assess the effects of DMRT1 loss on germ cell development, we monitored expression of a selected meiotic marker at E13.5 and E17.5, after the initiation of meiosis in the chicken (Figure 3a). Meiotic progression was assessed by monitoring γH2AX (gamma H2A histone family member X), an indicator of double-stranded DNA breaks^21,24^. As expected, this marker was not expressed in germ cells of Z^D+^Z^D+^ gonads at either developmental stage, while in germ cells in Z^D+^W gonads expressed γH2AX at both stages with a reduction at E17.5. In the germ cells of gonads from Z^D+^Z^D-^ embryos, γH2AX was present at both stages, although in E17.5 gonads, γH2AX expression was more abundant compared to Z^D+^W controls, indicating a potential delay in meiotic entry in Z^D+^Z^D-^ gonads. In the gonads of Z^D-^W embryos, there was no evidence of γH2AX expression at either developmental stage, suggesting a delay or failure of meiosis.

**Figure 3.**
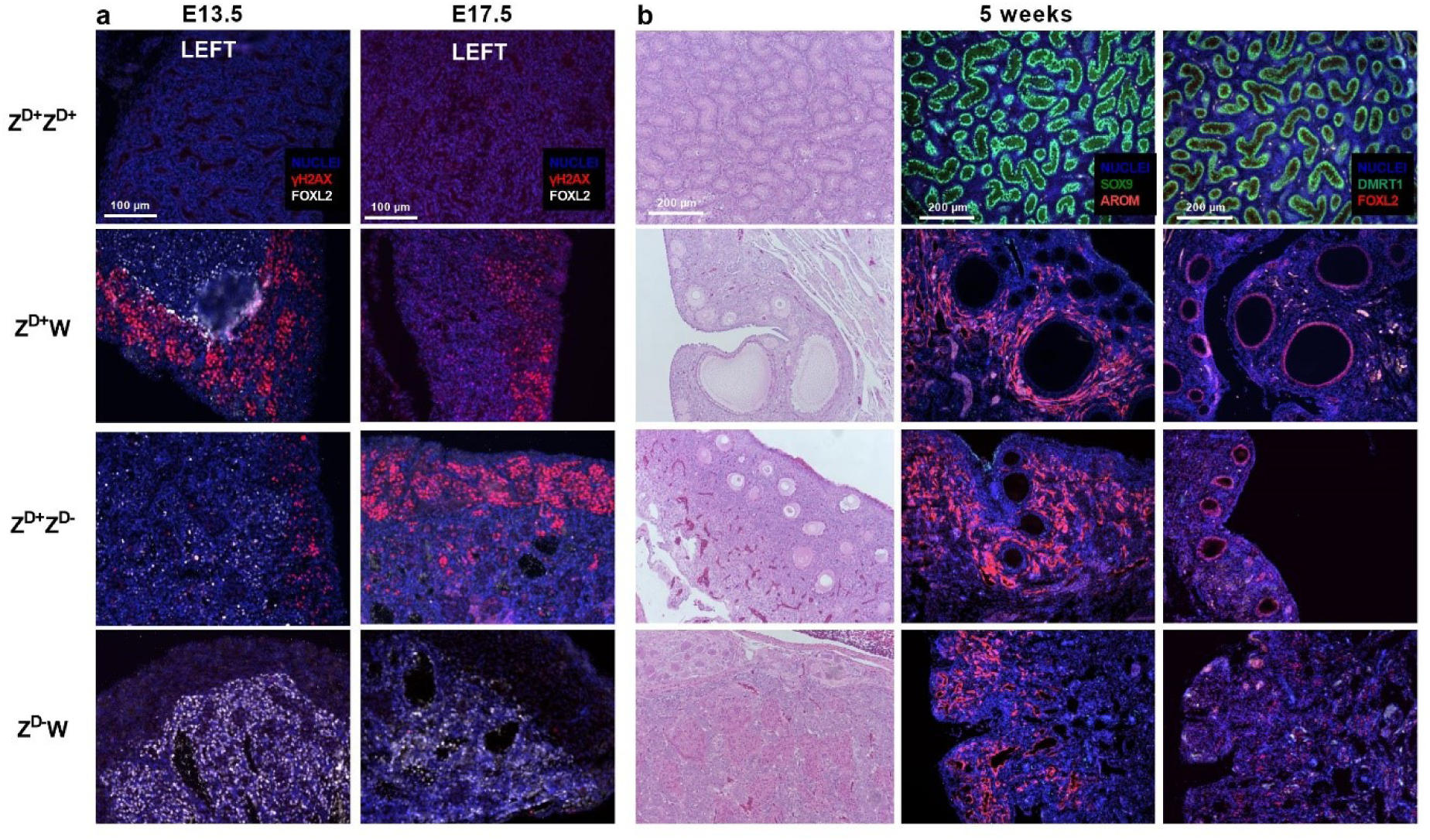
Effect of DMRT1 loss on follicular development. (a) FOXL2 and γH2AX expression in germ cells of gonads from wild type and *DMRT1*-mutant embryos at E13.5 and E17.5 of development. (b) Analysis of gonads of wild type and *DMRT1*-mutant birds at 5 weeks post-hatch. Sections were stained with either H&E or for testis or ovary-specific markers (FOXL2, AROM, SOX9 and DMRT1).

### Follicular development in *DMRT1*-mutant chicken

To determine whether the gonadal sex-reversal observed during embryonic development was permanent, we examined gonads of birds at five weeks post-hatch. Histological sections of gonads were stained with haematoxylin and eosin (H&E), or processed for IHC to examine expression of male and female markers (Figure 3b). The gonads of Z^D+^Z^D+^ birds exhibited typical testicular structures with seminiferous tubules showing strong expression of SOX9 and DMRT1. The gonads of Z^D+^W birds displayed a clear cortex with oocyte-containing follicles of different sizes. FOXL2 was highly expressed in the granulosa cells enclosing the oocyte, and aromatase was expressed in the thecal tissue surrounding the follicles. The structure and the expression patterns of FOXL2 and aromatase seen in the gonads of Z^D+^Z^D-^ birds was similar to the Z^D+^W birds and small follicles were clearly present. However, no larger follicles were observed in Z^D+^Z^D-^ birds. The gonads of Z^D-^W birds contained no oocytes/follicles and FOXL2 and aromatase were expressed in cells dispersed throughout the cortex. It is clear from this analysis that the testis-to-ovary sex-reversal in Z^D+^Z^D-^ birds was permanent and complete. It is well established that DMRT1 is highly expressed in both male and female germ cells and the absence of oocytes/follicles in the gonads of Z^D-^W birds, is likely a direct result of this leading to a perinatal failure of the germ cells to progress into meiosis. As expected, neither the Z^D+^Z^D-^ nor the Z^D-^W birds produced eggs (Supplementary Figure 4d).

### Gonadal sex-reversal does not affect secondary sex characteristics

We have previously established that chickens possess a degree of cell-autonomous sex identity (CASI) i.e. the secondary sexual phenotype depends, at least partly, on the sex-chromosome content of the somatic cells and not simply on gonadal hormones^17^. The generation of Z^D+^Z^D-^ birds that possess an ovary instead of testes enabled us to investigate the extent of CASI in chickens. In terms of secondary characteristics, male birds are heavier (possess greater muscle mass and bone density), they have larger combs and wattles, they possess hackle feathers (hood), and they develop leg spurs (Figure 4a). We assessed sexually mature adult birds at 24 weeks of age. It is clear from these images that the chromosomally male bird with an ovary (Z^D+^Z^D-^) was identical in appearance to the wild type Z^D+^Z^D+^ bird; with large comb and wattles, hackle feathers and obvious leg spurs. Z^D-^W birds were similar in appearance to Z^D+^W birds. Given that the Z^D+^Z^D-^ bird possesses an ovary rather than testes (Supplementary Figure 4d), this suggests that these typical male secondary sexual characteristics are due to CASI and independent of gonadal hormones.

**Figure 4.**
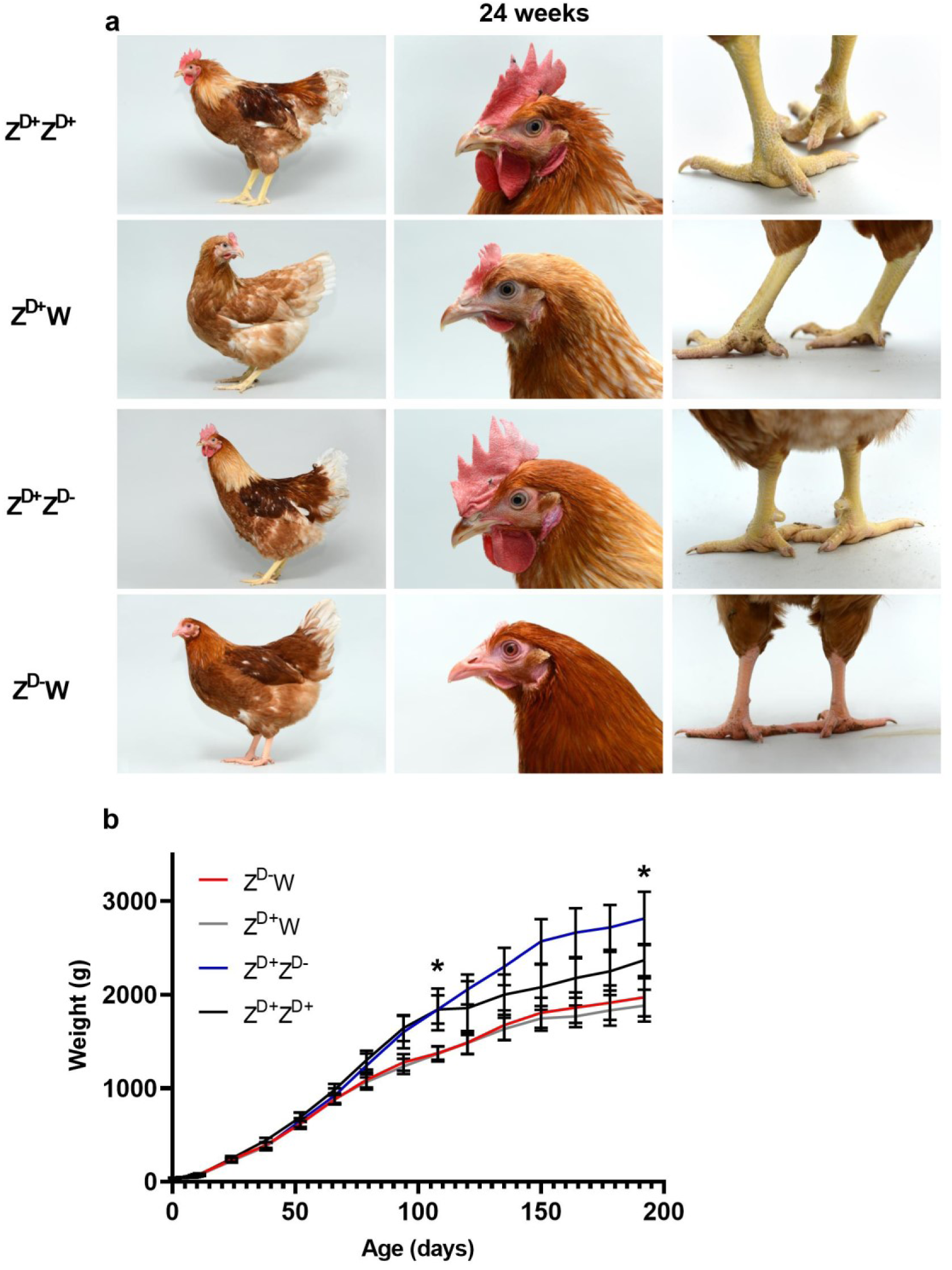
Phenotyping of adult *DMRT1* mutants. (a) Physical appearance of wild type and of *DMRT1*-mutant birds at 24 weeks. (b) Body weight of wild type and *DMRT1*-mutant birds. Asterisks indicate a statistically significant difference in body weight between each of the ZZ genotypes (Z^D+^Z^D-^, Z^D+^Z^D+^) and each of the ZW genotypes (Z^D+^W, Z^D-^W), on days 120 and 192.

We monitored the body weight of wild type and *DMRT1*-mutant birds over a 28-week period (Figure 4b). In this line of layer chickens, weights of wild type male and female birds diverge at 10 weeks (70 days), resulting in adult males that were approximately 20 % heavier than adult females. The Z^D-^W birds followed an almost identical growth pattern to Z^D+^W birds. Z^D+^Z^D-^ birds showed an identical weight increase to Z^D+^Z^D+^ birds up to 120 days, but then showed an even greater weight gain until 150 days of age. Post-mortem examination suggested that this additional weight accrues from abdominal fat deposits: a phenomenon also associated with capons^25^ (castrated cockerels; data not shown). These results suggest that the weight difference between the Z^D+^Z^D+^ birds and Z^D+^Z^D-^ was due to the loss of testes rather than the acquisition of an ovary. This further suggests that secondary sex characteristics of non-reproductive tissues in chickens are primarily due to the sex chromosome content of cells/tissues and independent of gonadal hormones.

Surprisingly, we observed that the Z^D+^Z^D-^ birds contained mature oviducts derived from both Mullerian ducts; in wild type male birds both Mullerian ducts regress, while in wild type female birds only the left Mullerian duct is retained, becoming the mature oviduct (Supplementary Figure 4b-c). In the adult Z^D+^Z^D-^ birds, two mature oviducts were present and connected to the cloaca. Examination of the reproductive ducts of E17.5 embryos showed that while the right Mullerian ducts of both Z^D+^W and Z^D-^W embryos had fully regressed, the right Mullerian ducts of Z^D+^Z^D-^ embryos exhibited only a slight shortening (Supplementary Figure 4a). It is well established that wild type female birds with one oviduct generate low levels of AMH during gonadal development, so the retention of both Mullerian ducts in Z^D+^Z^D-^ birds is consistent with a complete loss of AMH expression at embryonic stages (see Figure 2g and Supplementary Figure 3b).

### Female sex-reversal by E_2_-blockade requires DMRT1

Multiple reports have established that E_2_ plays a key role in ovarian differentiation in chickens ^10,26^. The epithelium of the left gonad, in both female and male embryos, expresses ERα, and this tissue responds to the presence of E_2_ by forming a thickened cortex containing germ cells. Studies with mixed-sex gonadal chimeras have shown that the presence of a small portion of aromatase-expressing ZW (ovarian) tissue is sufficient to induce cortex formation in the left gonad of wild type ZZ embryos^10^. It is also well established that blockade of the synthesis of E_2_ in Z^D+^W embryos, results in a sex reversal and the gonads develop as testes.

Here we assessed the effects of blocking E_2_ synthesis on gonadal development in *DMRT1* mutants: Z^D+^Z^D-^ and Z^D-^W. Fertile eggs carrying wild type and *DMRT1*-mutant embryos were injected with an inhibitor of aromatase activity (fadrozole) at E2.5 of development, and then re-incubated until E13.5 of development. Gonads were collected and processed for IHC. Sections of left gonads were stained for the presence of DMRT1 and for testicular and ovarian markers (Figure 5). The Z^D+^Z^D+^ gonad displayed obvious PGC-containing medullary sex cords with strong DMRT1 and SOX9 expression. The Z^D+^W gonad had a clear PGC-containing outer cortex and displayed medullary expression of FOXL2 and aromatase. The gonads of fadrozole-treated Z^D+^W embryos were clearly affected and showed clear evidence of female to male sex-reversal; the medulla contained sex cords with germ cells, aromatase expression was reduced and SOX9 expression was evident, and no cortex was present. Z^D+^Z^D-^ treated embryos displayed a similar pattern, demonstrating a rescue of the male to female sex reversal phenotype. This indicates that embryos with a single copy of DMRT1 will develop as testes in the absence of oestrogen. In contrast, fadrozole-treatment of Z^D-^W embryos did not result in female to male medullary sex-reversal; medullary sex cords did not form and the expression of FOXL2 and aromatase was maintained, however a thickened cortex is absent.

**Figure 5.**
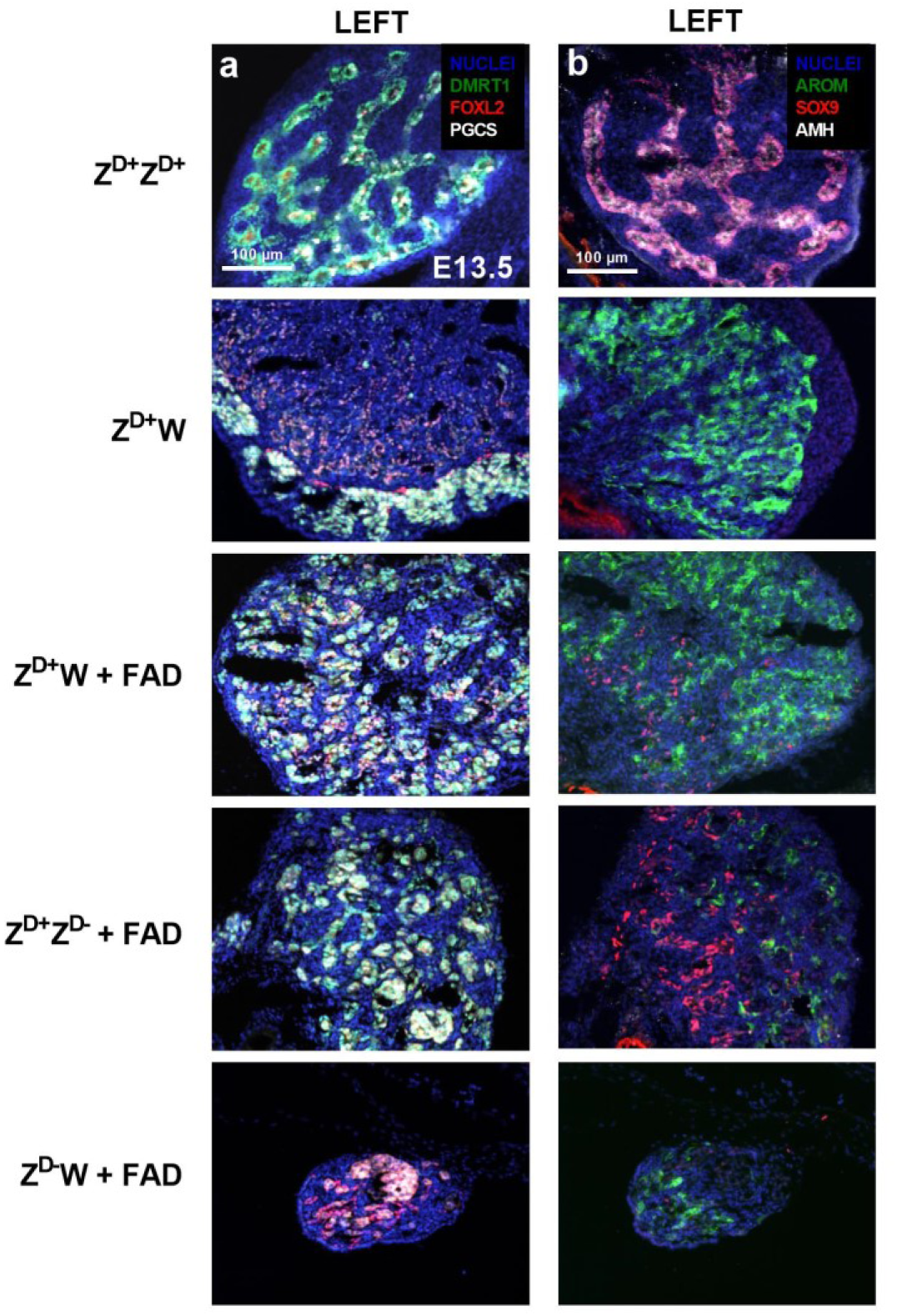
Expression of testis and ovary markers in gonads of fadrozole (FAD)-treated E13.5 embryos. Left gonads are shown. a) IHC of DMRT1, FOXL2 and PGC-marker (VASA), b) IHC of aromatase, SOX9 and AMH. FAD = Fadrozole-treated. Representative of three embryos per genotype.

These findings show that blocking E_2_ synthesis allows testis formation in Z^D+^Z^D-^, but not in Z^D-^ W embryos (Figure 5). Therefore, although a lack of E_2_ prevents the development of an obvious cortex in fadrozole-treated Z^D-^W embryos, DMRT1 is essential for testis development.

## Discussion

To clarify the role of DMRT1 in sex-determination and gonadal development in chickens, we used a CRISPR-Cas9 based approach to generate male offspring carrying disrupting mutations in *DMRT1*. Z^D+^Z^D-^ Genome edited PGCs were transmitted through a novel sterile surrogate host, leading to 100 % germline transmission. The G1 offspring presented the four chromosomal genotypes in a 1:1:1:1 ratio: Z^D+^Z^D+^, Z^D+^W, Z^D+^Z^D-^, and Z^D-^W. The equal transmission of all four possible genotypes demonstrates the Z^D+^ and Z^D-^ spermatozoa formed in the surrogate host gonad were all viable.

The gonads of Z^D+^Z^D-^ embryos resembled the gonads of wild type female embryos at the equivalent stage at all stages of development examined (E5.5 – E 17.5). These findings clearly demonstrate that the loss of a single copy of *DMRT1* in male birds results in ovarian rather than testicular development, and represent definitive proof of a DMRT1-dependent dosage-based mechanism of sex-determination in birds. To determine whether this switch in gonadal fate persisted post-hatch, we examined the gonads of these birds at five weeks of age, and again found that these resembled the gonads found in wild type females. The tissue is clearly ovarian with a thickened cortex containing follicles, with oocytes surrounded by granulosa and theca layers. Although these ovaries contained significant numbers of small and medium-sized follicles, there was a lack of large follicles and these birds did not ovulate/lay eggs at sexual maturity. In the wild type female (Z^D+^W), follicular maturation and ovulation is stimulated by signals from the hypothalamic-pituitary axis (HPA), and the lack of a female HPA in sex-reversed males (Z^D+^Z^D-^) may explain why follicles fail to mature.

Alternatively, this failure may be due to subtle defects in Z^D+^Z^D-^ granulosa or theca cells. In any event, the gonads of 5-week old Z^D+^Z^D-^ birds are clearly ovarian and demonstrate that the testis to ovary sex-reversal resulting from the loss of one functional copy of *DMRT1* is a permanent feature.

Unexpectedly, the *DMRT1* Z^D+^Z^D-^ birds were found to contain two mature oviducts. The right oviduct was shorter than the left oviduct, and in E17.5 embryos, the right Mullerian duct was also shorter than its left counterpart. The mechanism underlying persistence of the right Mullerian duct in Z^D+^Z^D-^ embryos is unclear, although regression in Z^D+^W embryos is thought to involve AMH or AMHR2 signalling. In any event, it appears that the retained mullarian duct tissue is able to respond to the same differentiation signals as the left Mullerian duct and generate a second oviduct. This was surprising, as a recent study concluded that DMRT1 was required for the early stages of Mullerian duct development ^27^. Our findings demonstrate that DMRT1 is not required for Mullerian duct development; the left Mullerian duct forms in Z^D-^W embryos that lack DMRT1 (Supplementary Figure 4d). It is possible that the different outcomes observed in these studies is due to differences in the timing of DMRT1 depletion. In our study, DMRT1 is absent throughout development, whereas in the earlier study, DMRT1 transcript levels were suppressed in the mesenchyme of the duct during elongation. Perhaps the early depletion of DMRT1 allows for the induction of a factor(s) that compensate for this loss and enable Mullerian duct formation.

We also analysed gonads of Z^D-^W embryos and found that loss of DMRT1 had little effect on gonadal sex identity, in that female embryos clearly had a left ovary with a thickened cortex containing germ cells. However, when we examined these ovaries at five weeks post-hatch, there were no obvious follicles and no evidence of oocytes, although the cortex did contain granulosa cells and theca cells. This suggests that the absence of functional DMRT1 leads to a loss of germ cells in post-hatch female birds. Given that *DMRT1* is highly expressed in germ cells and implicated in meiosis in other species, we analysed meiotic progression in late stage embryos (E13.5 & E17.5) by monitoring the expression of γH2AX. For Z^D+^Z^D-^ embryos, the pattern of marker expression in cortical PGCs was similar, although delayed, to that seen in wild type female embryos. In contrast, no γH2AX expression was detected in cortical PGCs of chromosomally female embryos lacking DMRT1 (Z^D-^W): a similar PGC phenotype to that observed in *DDX4*-mutant chickens, where the germ cells are lost^21^. Taken together, these findings suggest that in these birds the loss of DMRT1 either prevented or delayed meiosis and resulted in the loss of germ cells.

It is clear from our studies that the loss of one copy of *DMRT1* in chromosomally male embryos results in the induction of the gene network underlying ovary development: the spatial and temporal expression of first FOXL2 and then aromatase is identical to that seen in wild type female embryos. This suggests that the presence of two functional copies of *DMRT1* in wild type male embryos suppresses, either directly or indirectly, the expression of FOXL2. In goats, FOXL2 is a primary ovarian determinant; it has been shown to be a direct activator of aromatase, which catalyses the conversion of androgens to oestrogen^28-30^. It is well established that E_2_ also plays a major role in sex-determination in birds. Oestrogen treatment of chromosomally male embryos leads to ovary formation and inhibition of E_2_ synthesis in chromosomally female embryos results in ovary to testes sex-reversal^8,10^. In this study, we have investigated the effects of blocking E_2_ synthesis in embryos with targeted mutations in *DMRT1*. We have demonstrated that the left gonad in Z^D+^Z^D-^ embryos develops as an ovary, however, if E_2_ synthesis is blocked in these embryos, both gonads develop as testes. Interestingly, when E_2_ synthesis is blocked in chromosomally female embryos that lack DMRT1, the gonads do not develop as testes, suggesting that DMRT1 is essential for testis formation. The gonad medulla of these embryos continues to express FOXL2 and aromatase, but because E_2_ synthesis is blocked, cortex formation is not induced. It is noteworthy that the early gonads of Z^D-^W embryos are smaller than those of Z^D+^W embryos, perhaps reflecting a requirement for DMRT1 in the cellular allocation and/or proliferation of the early gonad. Figure 6a summarises the fate of the gonadal medulla and cortex under the influence of different combinations of DMRT1 and E_2_. We hypothesise that primary sex-determination in chickens depends on whether or not the gonadal medulla expresses E_2_. In Z^D+^Z^D+^ embryos, high levels of the Z chromosome DMRT1 suppress FOXL2 expression, which in turn leads to an absence of aromatase and to low levels of E_2_ synthesis and allows sex cord formation to be induced. In Z^D+^W embryos, levels of DMRT1 are not sufficient to suppress FOXL2 and the resulting E_2_ inhibits the testis network and induces cortex formation. If E_2_ synthesis is blocked in Z^D+^W embryos, or Z^D+^Z^D-^ embryos, the male pathway is not inhibited and testis development occurs. If E_2_ synthesis is blocked in embryos devoid of DMRT1 (Z^D-^W), the medulla develops an ovarian phenotype, suggesting that DMRT1 is required for testis formation and PGC survival, but it is not necessary for ovary development.

**Figure 6.**
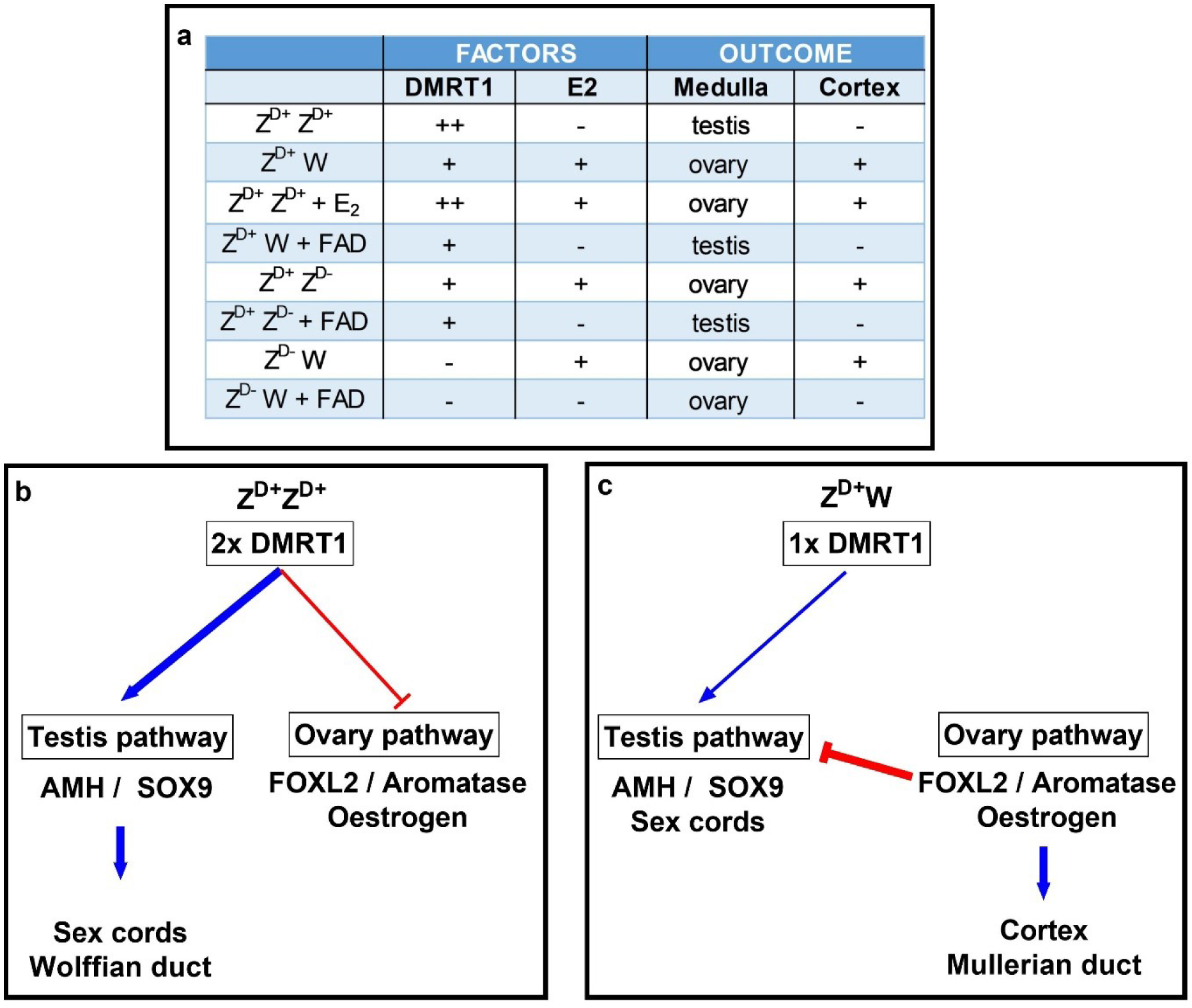
Overview of sex-determination in chickens. a) Outcomes resulting from different combinations of DMRT1 and E_2_. b, c) Schematics illustrating regulation of gene networks that define male and female reproductive systems (DMRT1: ++ / + / - = 2 / 1 / 0 copies; E_2_ and Cortex: +/ - = present/ absent).

Previously it was considered that the male and female secondary sexual characteristics of vertebrates were largely dependent on the outcome of primary sex-determination, and that gonadal hormones played a major role in defining the sexual phenotype. More recently it has become generally accepted that male:female differences are due to the combined effects of gonadal hormone differences and differences in the sex-chromosome constitution of individual cells and tissues, a classic example being that of marsupial body dimorphism (reviewed here^31^). We and others have established that birds possess a cell autonomous sex identity (CASI) and that this plays a major role in defining secondary sexual characteristics^17,32,33^. Analysis of the adult birds in this study suggest that CASI may be the dominant factor in establishing sexual phenotype and that gonadal hormones have little or no effect on external secondary sexual characteristics The male birds with ovary in place of testes are virtually identical in growth rate and appearance to wild type males and display no female characteristics.

Taken together, our findings clearly place DMRT1-dosage in the centre of the avian gonadal sex determining mechanism, while providing evidence for an important role of DMRT1 in germ cell and Mullerian ducts fate. Finally, this work further highlights the unique feature of cell-autonomous sex identity in birds.

## Methods

### Genome editing and generation of *DMRT1* mutant birds

Germ cells were isolated from Hy-line Brown layer embryos heterozygote for an RFP reporter gene^34^ at HH stage 16^+^ (Hamburger & Hamilton) and cultured *in vitro*^*35*^. Briefly, 1 μl of embryonic blood was aspirated from the dorsal aorta of embryos and placed in FAOT culture medium^36^. Expanded germ cell populations (3 weeks) were co-transfected with 1.5 μg of high fidelity CRISPR-Cas9 vector (HF-PX459 V2.0) which included a targeting guide (sgRNA) for the *DMRT1* locus and two single-stranded donor oligonucleotides (ssODNs, 5 pmol of each, see Supplementary Table 1) using Lipofectamine 2000 (Thermo Fisher Scientific, ^20^). Twenty-four hours after transfection, PGCs were treated with Puromycin (at 400 ng/mL) for 48 hours to select for edited cells. Following puromycin treatment, PGCs were sorted into single wells of 96-well plates using a FACSAria III (BD Biosciences) at one PGC per well in 110 μL FAOT to produce clonal populations. PGCs were expanded in culture, DNA was extracted for analysis, and then clonal PGCs were cryopreserved in STEM-CELLBANKER (AMSBIO).

### Generating Surrogate Host Chicken

Clonal PGCs were thawed and 1 μl of cells from an individual PGC clone carrying the desired edits for DMRT1 were injected via the dorsal aorta into stage 16 HH+ transgenic surrogate host embryos containing an inducible Caspase9 targeted to the germ cell-specific *DAZL* locus (Ballantyne et al, under review; ^37^). 1.0 μl of 25mM B/B (in DMSO) (AP20187, Takara) was added to 50ul of PGCs (3,000 PGCs/μl) before injection and subsequently 100ul P/S (containing 3ul of 0.5mM B/B drug (in EtOH) was pipetted on top of the embryo. Treatment of the transgenic surrogate hosts with B/B drug ablates the endogenous germ cells, such that the only gametes that can form are from the donor PGCs. Fourteen surrogate host chicks were hatched from two injection experiments. Four surrogate host chicks carried the iCapsase9 transgene. Two male iCaspase9 surrogate hosts carrying germ cells heterozygous for DMRT1 (Z^D+^Z^D-^) were crossed with wild type hens (Z^D+^W) to produce G1 embryos for analysis and hatched to create G1 offspring. All animal experiments were conducted under UK Home Office licence.

### Genetic screening

DNA was extracted from cells and embryonic tissues using the PureLink Genomic DNA Mini Kit (Thermo Fisher Scientific) according to the manufacturer’s instructions. To amplify the *DMRT1* locus, PCR reactions included 100ng gDNA, and Q5 high-fidelity polymerase (New England Biolabs) and comprised the following cycling parameters: 98°C for 2min, 98°C for 30s, 68°C for 30s, 72°C for 30s, 72°C for 2min (steps 2 to 4 run for 32 cycles; Forward primer: CATGCCCGGTGACTCCC; Reverse primer: GATCAGGCTGCACTTCTTGC). Gene editing included insertion of a Hindlll restriction site, and to screen clones PCR products were digested using HF-HindIII (NEB). Enzyme digests were separated by electrophoresis and genotypes distinguished by fragment banding patterns (wild type, mono-allelic and bi-allelic *DMRT1* mutants, Supplementary Figure 1). All PGC cultures and chick embryos were sexed using a rapid, invader-based sexing assay ^38^.

### Tissue collection

Freshly laid fertile eggs were incubated blunt side up, at 37.5°C, in 60 % humidity, with rocking (one rotation per 30 minutes) for the desired incubation period.

Eggs were removed from the incubator at the required stage (E5.5, E6.5, E8.5, E13.5 and E17.5) and embryos were carefully removed, sacrificed according to Home Office Schedule I procedures and the gonads dissected and processed for further analysis. Gross morphology of gonads was recorded using a Zeiss Axiozoom Microscope (Carl Zeiss AG).

For RNA analysis, gonads were dissected, placed in PBS, and any remaining of mesonephric tissue removed. Gonads were snap-frozen in 10 μL of RNA-Bee (AMS Biotechnology) until RNA extraction. For Western analyses, gonads were collected into 100 μL of RIPA buffer (Thermo Fisher Scientific). For immunostaining, gonads+mesonephroi were placed in 4 % paraformaldehyde (see below). A small portion of embryonic wing tissue was collected and used to determine genetic sex.

### Quantitative Real Time PCR

Individual gonad pairs from E8.5 embryos were homogenized in RNA-bee (AMS Biotechnology) and the lysate was loaded onto a Direct-zol RNA Microprep RNA extraction column (Zymo Research) and DNase-treated as per the manufacturer’s protocol. First-strand cDNA was synthesized using the ‘First-strand cDNA synthesis kit’ (GE Healthcare) according to the manufacturer’s instructions. Primers were designed to amplify transcripts from the following genes: *DMRT1, FOXL2, AROM, SOX9, and AMH*. PCR reactions were optimised to meet efficiencies of between 95 % and 105 % across at least a 100-fold dilution series (primer sequences are listed in Supplementary Table 1). QPCR reactions were performed using a Stratagene MX3000P qPCR system (Agilent Technologies). The chicken hydroxymethylbilane synthase gene (HMBS) was used as an internal control^39^. Data were analysed using the 2^-ΔΔCt^ method^40^.

### Western blotting

Gonads were collected in RIPA buffer (Thermo Fisher Scientific) and disrupted with a handheld homogeniser. Protein levels were quantified using a Pierce BCA protein assay kit (Thermo Fisher Scientific). Protein samples (10 μg) were separated on 4 % - 15 % Bis-tris gels (Bio-Rad Laboratories) and wet-transferred onto a PVDF membrane. Membranes were blocked in Intercept Blocking Buffer for 1 hour (LI-COR Biosciences) and incubated overnight with primary antibodies; rabbit anti-DMRT1^41^, rabbit anti-γ-tubulin, T3559, Sigma. After four washes in TBST, blots were incubated with secondary antibody (HRP-conjugated) for 1 hour at room temperature, followed by four washes in TBST. Hybridisation signals were detected using of a Novex chemiluminescence kit (Life technologies) and membranes exposed to Hyperfilm ECL (Amersham). Membranes were stripped for 10 minutes in Restore PLUS Western Blot stripping buffer (Thermo Scientific) for re-hybridisation.

### Immunohistochemistry

Immunohistochemistry was carried out according to the protocol described by Stern^42^. Gonads were fixed in 4 % paraformaldehyde for 2 hours at 4°C. Tissues were equilibrated in 15 % sucrose/0.012M phosphate buffer overnight, embedded in 15 % sucrose plus 7.5 % gelatin/0.012M phosphate buffer (pH 7.2) and snap frozen using isopentane. Ten micrometer (10 μm) thick sections were cut on a cryostat (OTF 5000 Bright Instruments) and mounted on Superfrost Plus slides (Thermo Fisher Scientific). Slides were de-gelatinised for 30 min in PBS at 37°C and blocked in PBS containing 10 % donkey serum, 1 % BSA and 0.3 % Triton X-100 for 2 hours at room temperature. Incubation with primary antibodies (Supplementary Table 2) was carried out overnight at 4°C, followed by washing four times in PBS containing 0.3 % Triton X-100, and incubation with secondary antibodies for 2 hours at room temperature. After washing four times in PBS containing 0.3 % Triton X-100, the sections were treated with Hoechst nuclear stain solution (10 μg/ml) for 5 min. Imaging was carried out using a Leica DMLB Upright Fluorescent microscope (Leica Camera AG).

### Data analysis

All summary data values are expressed as mean ± standard deviation. GraphPad Prism (Graphpad) was used to produce graphs and for statistical analyses. Statistical analysis of qPCR data included a one-way ANOVA analysis followed by Tukey’s multiple comparison test for *post-hoc* comparisons. P < 0.05 was set as the statistical significance threshold.

## Acknowledgements

Funding for this work was from the Biotechnology and Biological Sciences Research Council (BBSRC, BB/N018672/1), The Roslin Institute ISP funding grants, The Francis Crick Institute core funding to R.L.-B., which includes Cancer Research UK (FC001107), the UK Medical Research Council (FC001107) and the Wellcome Trust (FC001107); the UK Medical Research Council (U117512772 to R.L.-B.). The authors are grateful to Prof. M.A. Hattori for kindly providing VASA antibody. We wish to thank the National Avian Research Facility at the Roslin Institute for animal husbandry services and the Bio-imaging Facility at the Roslin Institute for technical assistance.

## Footnotes

### Author Contributions

Conceptualization: M.C., M.M., S.G., R.L.-B.; Methodology: J.I., G.T., D.Z., L.L., A.I.-A.; Validation: J.I., D.Z.; Formal analysis: J.I., D.Z. M.C., M.M.; Investigation: J.I., G.T., D.Z., L.L.; Writing - original draft: J.I.; Writing - review & editing: J.I., G.T., D.Z., L.L., D.G., A.I.-A., R.L.-B., S.G., M.M., M.C.; Visualization: J.I.; Project administration: M.C., R.L.-B., S.G., M.M.; Funding acquisition: M.C., R.L.-B., M.M., S.G.

