## Supplementary Material for "Primary sex determination in chickens depends on DMRT1 dosage, but gonadal sex does not determine secondary sexual characteristics in adult birds"

**Supplementary materials**

**Supplementary Table 1:** Sequences of primers used for RT-qPCR and DMRT1 CRISPR ssODNs.

| Target | Forward | Reverse |
|---|---|---|
| *DMRT1* | TTCGCGTTGAGTGCCTCGAC | TTCGCGTTGAGTGCCTCGAC |
| *FOXL2* | ACATGTTCGAGAAGGGCAAC | TGTTCATGAAGGTGGACTGC |
| *SOX9* | TACGATTACACCGAGCACCA | GTAGACGGGCTGTTCCCAGT |
| *AMH* | TCCACAGCTTTGCCGAGAA | CACATTTTCCCTCTTTGCTCCA |
| *AROM* | GCTATTTCCAGCCATTTGGA | TTGGGTGCATGGATAAGTCA |
| Mutant ssODN | CCCGCTGCCGCAACCACGGCTACTCCTCGGAATTCTAGGTCACAAGCGGTTCTGCTAGTGGCGGGACTGCCAGTGCAAGAAGTGCAGCCTGATC | |
| Synonymous ssODN | CCCGCTGCCGCAACCACGGCTACTCCTCGCCGCTTAAGGGTCACAAGCGGTTCTGCATGTGGCGGGACTGCCAGTGCAAGAAGTGCAGCCTGATC | |

**Supplementary Table 2:** Details of antibodies used for IHC

| Target | Details |
|---|---|
| DMRT1 | Rabbit, in-house, 1:500[1] |
| FOXL2 | Goat, Abcam, ab5096, 1:250 |
| SOX9 | Rabbit, AbD Serotec, AB5535, 1:250 |
| AMH | Goat, SCBT,SC-6886, 1:250 |
| P450 (AROM) | Mouse, AbD Serotec, MCA20775, 1:250 |
| PGC markers | VASA, Rat, 1:1000[2]<br>DAZL, Abcam, ab215718, 1:500 |
| γH2AX | Mouse, Merck, 05-636, 1:250 |
| γ-tubulin | Mouse ,Sigma, T5326, 1:1000 |

**Supplementary Table 3:** Genotype analysis of *DMRT1*-mutant embryos collected throughout study.

| | $Z^{D+}Z^{D+}$ | $Z^{D+}W$ | $Z^{D+}Z^{D-}$ | $Z^{D-}W$ | Total |
|---|---|---|---|---|---|
| **No. of embryos** | 53 | 43 | 56 | 51 | 203 |
| **Percentage** | 26.1% | 21.2% | 27.6% | 25.1% | 100.0% |

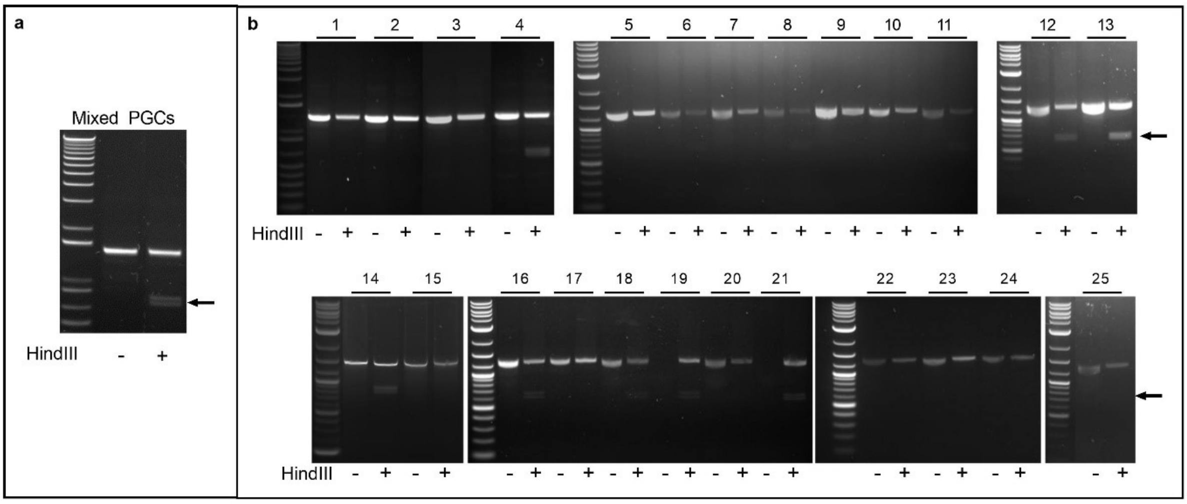


**Supplementary Figure 1:** Restriction analysis of *DMRT1*-edited PGCs. *Hind*III enzyme was used to analyse the *DMRT1* locus for the 'null' *DMRT1* allele (see Fig 1). Digested *DMRT1* PCR products from puromycin-selected mixed PGCs (a) and clonally cultured PGCs (b). *DMRT1* PCR product: 1.3kb; Restriction fragments: 0.6 kb and 0.7 kb

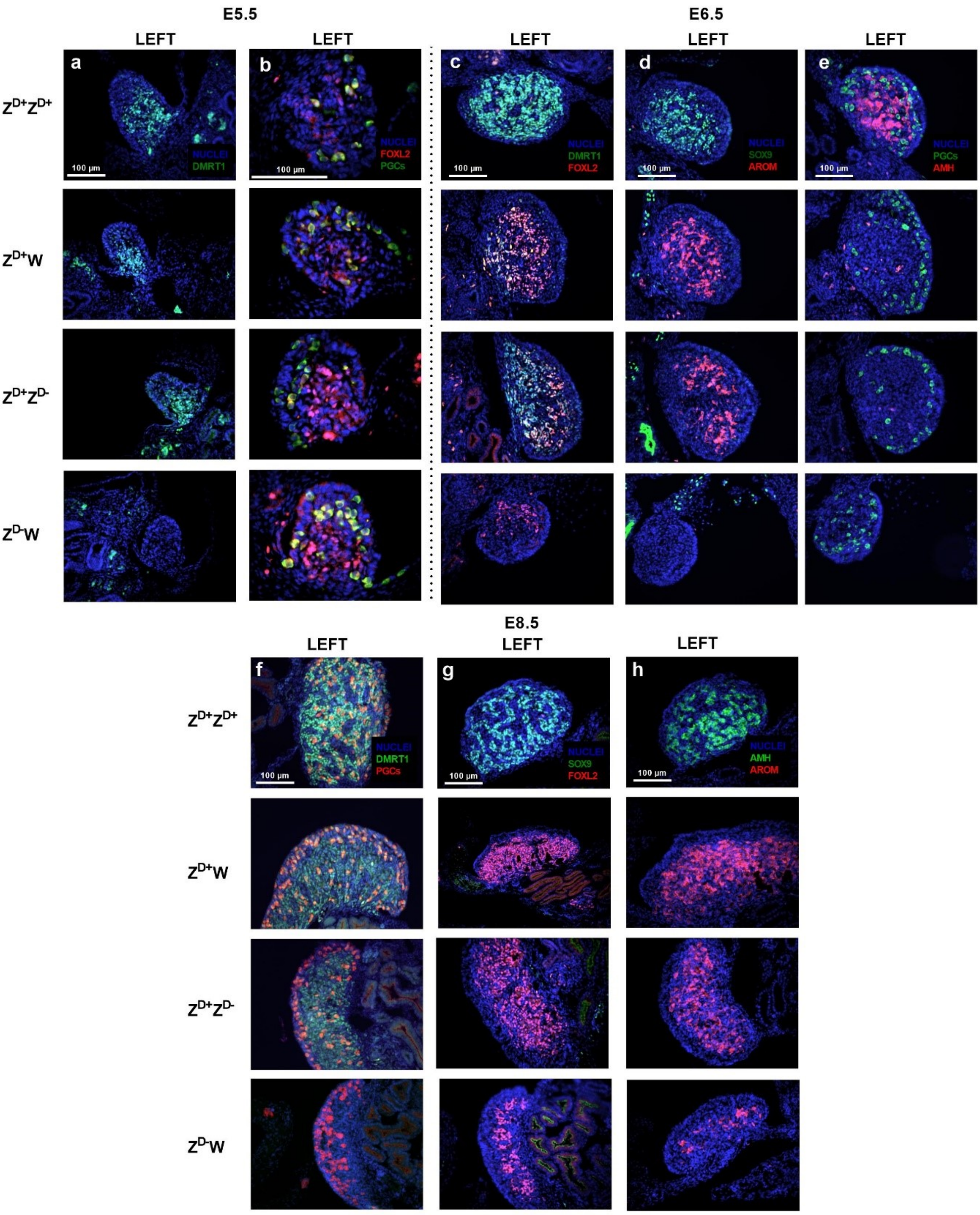


**Supplementary Figure 2**: Immunostaining of gonadal sections from E5.5 (a-b), E6.5 (c-e) and E8.5 (f-h) *DMRT1*-mutant embryos. Antibodies were against DMRT1, FOXL2, SOX9, AMH, aromatase (AROM) and germ cell markers (DAZL at E5.5 and E6.5, VASA at E8.5). Aromatase, SOX9 and AMH were not detected in E5.5 sections. There was no staining for aromatase in the $Z^{D-}W$ embryos on E6.5, suggesting a delay in initiating transcription. A minimum of three embryos of each genotype were examined.

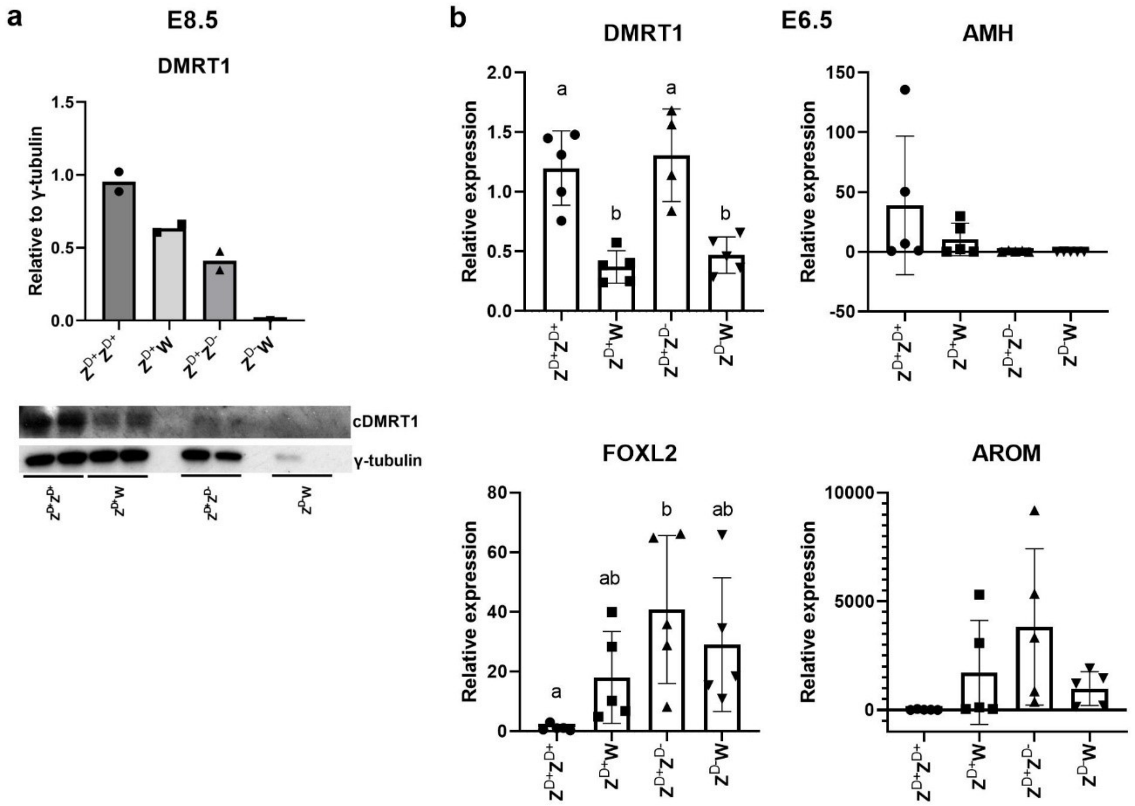


**Supplementary Figure 3:** Western blots (a) on *DMRT1*-mutant and wild type control gonads at E8.5. Histograms represent the mean intensity value derived from two lanes per genotype (with the exception of $Z^{D-}W$). Approximate band sizes for DMRT1: 45 kDa, γ-tubulin: 48 kDa. RT-qPCR profiling (b) of key gonadal markers in gonads from E6.5 *DMRT1*-mutant embryos (pool of five gonad pairs per data point). The degree of variation in expression seen within each genotype is likely to reflect slight differences in stage of development: *AMH*, *FOXL2* and *Aromatase* (AROM) sex-biased expression initiates around this stage. Note the low expression of *DMRT1* transcript in $Z^{D-}W$ gonads; this is attributed to the *DMRT1* primers detecting mutant *DMRT1* transcript, which is still produced (however, the mutant stop codon prevents translation of wild type protein). Bars represent the mean ± standard deviation relative to $Z^{D+}Z^{D+}$. Different letters signify statistically different expression between individual genotypes (see Methods).

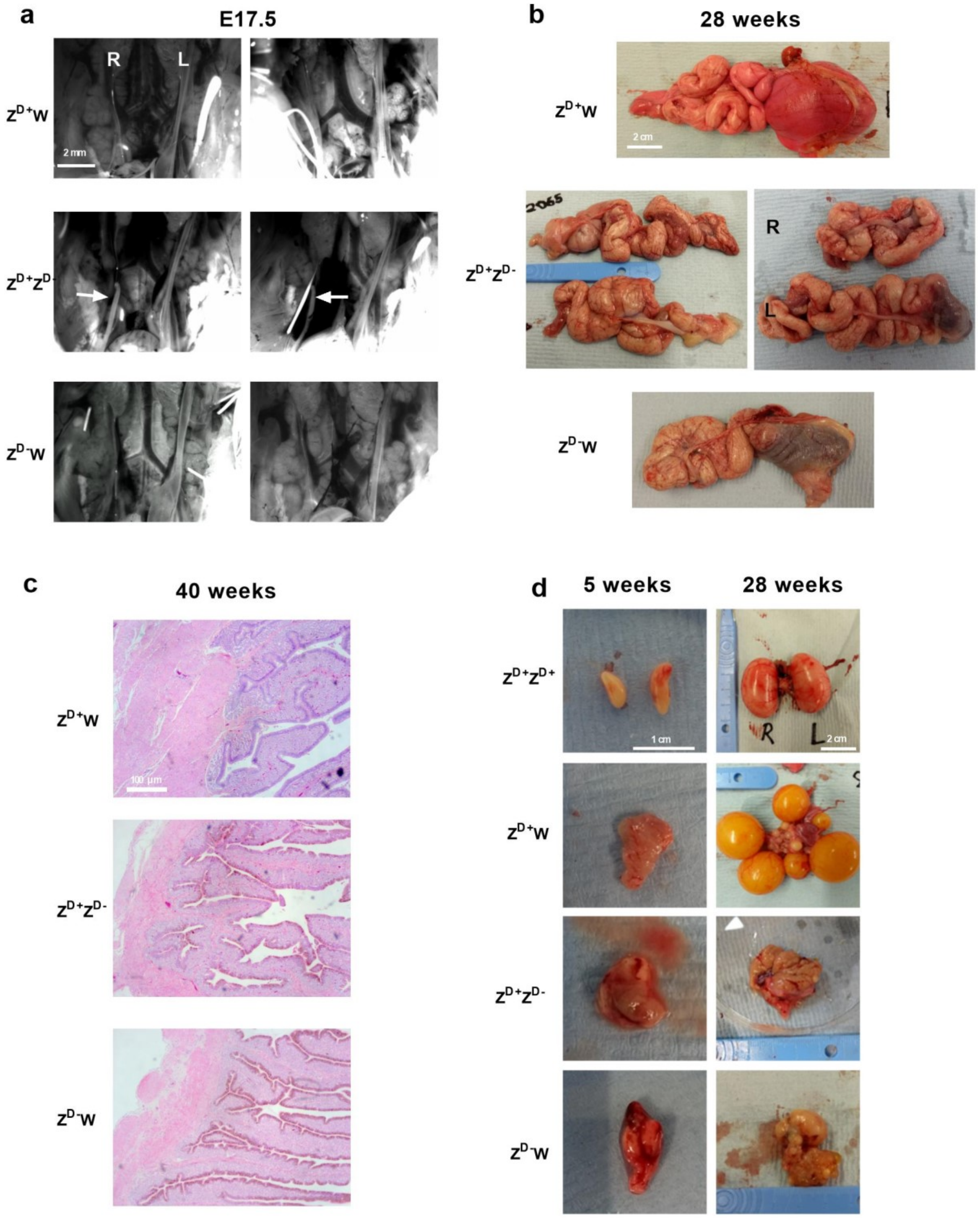


**Supplementary Figure 4:** Gross reproductive morphology of Mullerian ducts and gonads of *DMRT1* mutants. Images of Mullerian ducts (a) and oviducts (b) from *DMRT1* mutants and wild type controls at E17.5 and 28 weeks, respectively. c) Histological slides (H&E) of oviducts from *DMRT1* mutants and wild type controls at 40 to 45 weeks. Note the pigmentation in the epithelium of the non-laying genotypes $Z^{D+}Z^{D-}$ and $Z^{D-}W$. No histological abnormalities noted for the oviduct tissues examined. d) Photographs of *DMRT1*-mutant gonads and wild type controls at 5 weeks and 28 weeks post-hatch. Arrows in (a): retained right Mullerian ducts.
